# Free spermidine evokes superoxide radicals that manifest toxicity

**DOI:** 10.1101/2021.09.04.458997

**Authors:** Vineet Kumar, Rajesh Kumar Mishra, Debarghya Ghose, Arunima Kalita, Anand Prakash, Gopa Mitra, Amit Arora, Dipak Dutta

**Author notes:** Address correspondence to Dipak Dutta. Regional Centre for Biotechnology, Faridabad, Haryana, India (Vineet Kumar). Department of Medical Microbiology, Post Graduate Institute of Medical Education and Research, Research Block A, Sector-12, Chandigarh-160012, India (Amit Arora). Vineet Kumar and Rajesh Kumar Mishra contributed equally to this work.

## Abstract

Spermidine and other polyamines alleviate oxidative stress, yet excess spermidine seems toxic to *Escherichia coli* unless it is neutralized by SpeG, an enzyme for the spermidine N-acetyl transferase function. Besides, a specific mechanism of SpeG function conferring pathogenic fitness to *Staphylococcus aureus* USA300 strain is unknown. Here, we provide evidence that although spermidine mitigates oxidative stress by lowering hydroxyl radical and hydrogen peroxide levels, excess of it simultaneously triggers the production of superoxide radicals, thereby causing toxicity in the *E. coli* Δ*speG* strain as well as naturally *SpeG-*deficient *S. aureus* RN4220 strain. However, wild-type *E. coli* and *S. aureus* USA300 with a horizontally-acquired *speG* gene tolerate applied exogenous spermidine stress. Furthermore, we demonstrate that while RNA-bound spermidine inhibits iron oxidation, free spermidine interacts and oxidizes the iron to evoke superoxide radicals directly. Superoxide radicals thus generated, further affects redox balance and iron homeostasis. Since iron acquisition and metabolism in the host tissues is a challenging task for *S. aureus*, the current findings helped us explain that the evolutionary gain of SpeG function by *S. aureus* USA300 strain allows it to neutralize exogenous spermidine- and spermine-mediated toxicity towards iron metabolism inside the host tissues.

## Introduction

Polyamines are ubiquitously present in all life forms. They tweak a diverse array of biological processes, e.g., nucleic acid and protein metabolism, ion channel functions, cell growth and differentiation, mitochondrial function, autophagy and aging, protection from oxidative damage, actin polymerization, and perhaps many more (Casero et al., 2018; Gawlitta et al., 1981; Madeo et al., 2018; Michael, 2018; Miller-Fleming et al., 2015; Oriol-Audit, 1978; Pegg, 2016; Pegg, 2018; Pohjanpelto et al., 1981; Tabor and Tabor, 1984; Wallace et al., 2003). The cationic amine groups of polyamines can avidly bind to the negatively charged molecules, such as RNA, DNA, phospholipids, etc. (Igarashi et al., 2000; Miyamoto et al., 1993; Schuber, 1989; Tabor and Tabor, 1984). Polyamines have been demonstrated to protect DNA from reactive oxygen species (ROS) such as singlet oxygen, hydroxyl radical (•OH), or hydrogen peroxide (H_2_O_2_) (Balasundaram et al., 1993; Ha et al., 1998a; Ha et al., 1998b; Jung and Kim, 2003; Khan et al., 1992a; Khan et al., 1992b; LØVaas, 1996; Pegg, 2018; Stewart et al., 2018). Indeed, knocking out polyamine biosynthesis enzymes from *E. coli* and yeast confers toxicity to oxygen, superoxide anion radical (O_2_**^-^),** and H_2_O_2_ (Balasundaram et al., 1993; Chattopadhyay et al., 2003; Chattopadhyay et al., 2006; Eisenberg et al., 2009).

Most prokaryotes, including *E. coli* synthesize cadaverine, putrescine, and spermidine, while higher eukaryotes additionally synthesize spermine. *E. coli* also acquires spermidine and putrescine from the surrounding medium (Igarashi and Kashiwagi, 2000; Miller-Fleming et al., 2015). However, polyamine in excess is toxic to the organisms unless polyamine homeostasis in the cell is operated at the levels of export, synthesis, inactivation, and degradation (Miller-Fleming et al., 2015). Notably, spermine/spermidine N-acetyl transferase (SSAT or SpeG), which inactivates spermidine and spermine, constitutes the most potent polyamine homeostasis component of the cells (Miller-Fleming et al., 2015). Interestingly, *S. aureus* neither synthesizes polyamine nor encodes the SpeG enzyme. Thus, *S. aureus* strains are inherently hypersensitive to spermidine (Joshi, 2012). However, *S. aureus* USA-300 lineage, which is evolved through acquiring *speG* gene nested in the Arginine Catabolic Mobile Element (ACME), is exceptionally pathogenic (Joshi, 2012). The mechanism behind such a phenomenon is entirely unknown.

A tremendous volume of work has been dedicated to unravel the biological importance of spermidine and its homeostasis mechanisms. It has also been known for long that spermidine (or spermine) in excess is toxic to the organisms and viruses (Pegg, 2013). It has been proposed that the excess polyamines may affect protein synthesis by binding to acidic sites in macromolecules, such as nucleic acids, proteins and membrane, and by displacing magnesium from these sites (Pegg, 2013; Limsuwun and Jones, 2000). However, a precise molecular detail of spermidine toxicity is not yet understood. In this study, we decipher a unique molecular mechanism of spermidine toxicity in bacteria. We find the intertwined relationships among spermidine toxicity, iron metabolism, and O_2_**^-^** radical production in bacteria.

## Results

### Spermidine stimulates O_2_^-^ production while inhibits •OH and H_2_O_2_ production in *E. coli*

The optimum level of intracellular spermidine in wild-type (WT) *E. coli* is about 7 mM (Igarashi and Kashiwagi, 2000; Miyamoto et al., 1993). However, this level is expected to be elevated in the Δ*speG* strain or when administered exogenously. To determine the working concentrations of exogenous spermidine that sufficiently inhibits the growth of Δ*speG*, but not WT strain, we added various amounts of spermidine in the growth medium. WT cells showed a modest reduction in growth up to 6.4 mM of spermidine concentration (Figure 1 - figure supplement 1). On the contrary, Δ*speG* strain exhibited a striking decrease in growth when supplemental spermidine level was > 3.2 mM (Figure 1 - figure supplement 1). Therefore, we chose spermidine concentration ≥ 3.2 mM for our further experiments.

**Figure 1.**
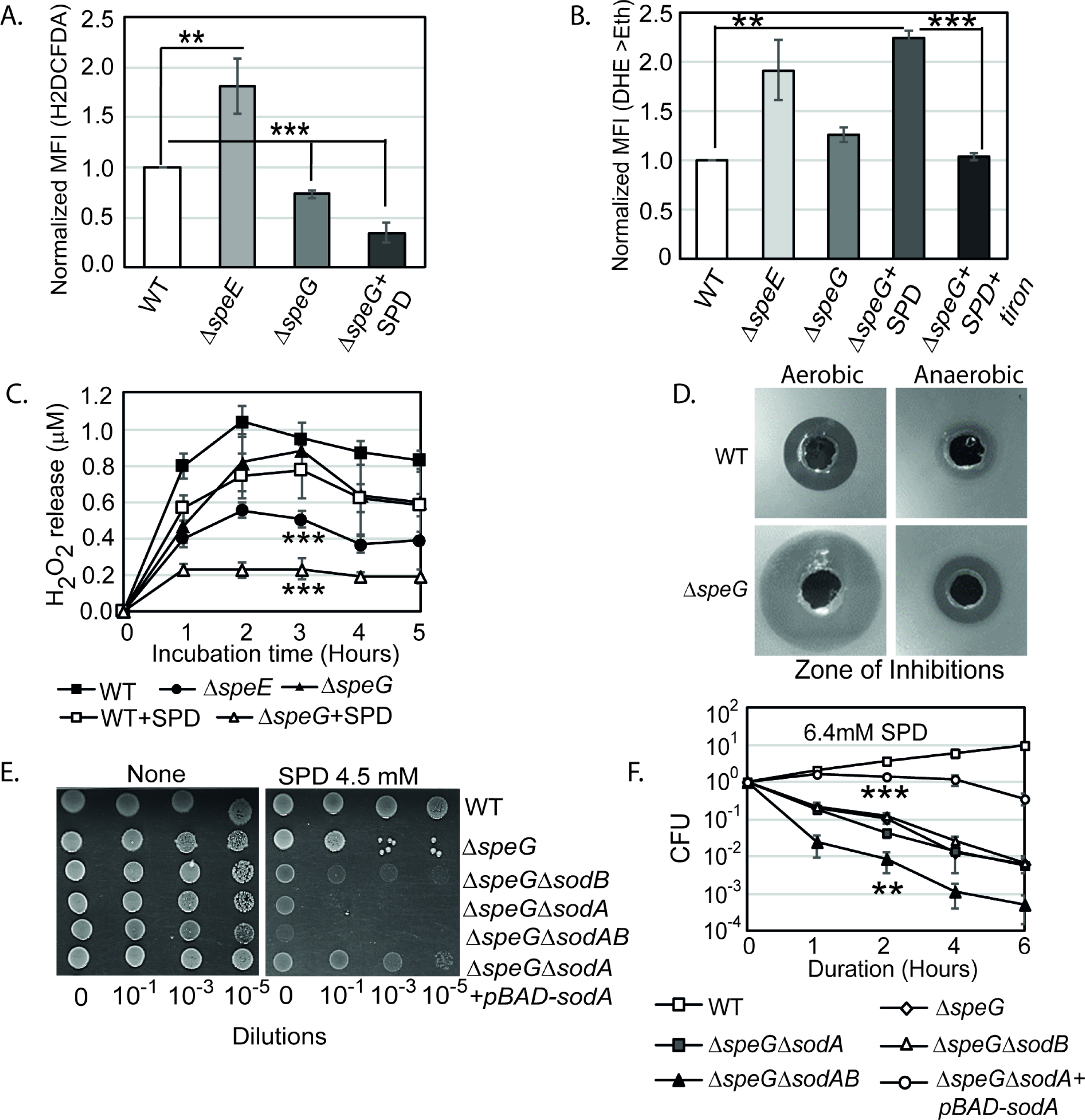
Spermidine (SPD) stress and intracellular ROS in *E. coli*. **A.** The relative MFI values for the H2DCFDA, which is an indicator of •OH radical production, obtained by flow cytometry analyses are plotted. **B.** The relative MFI values of DHE probe, which is an indicator of O_2_**^-^** radical production, obtained by flow cytometry analyses are plotted. **C.** The absolute H_2_O_2_ production for a span of 5 hours from the different *E. coli* strains are shown. *** are P values generated comparing with WT value. **D.** ZOIs surrounding SPD well on the agar plates were shown for the WT and Δ*speG* strains of *E. coli* under aerobic and anaerobic conditions. **E.** Serially diluted *E. coli* cells were spotted on LB-agar plates to show their sensitivity to SPD. **F.** Viability of different knockout strains were plotted from the CFU counts in different time intervals after treatment with lethal dose of SPD. ** and *** are P values generated comparing with the values of Δ*speG* and Δ*speG*Δ*sodA*, respectively. Error bars in the panels are mean ± SD from the three independent experiments. Whenever mentioned, *** and ** are <0.001 and <0.01, respectively; unpaired T test. See also Figure 1 – figure supplement 1, and source data 1-4.

Although spermidine is generally considered an anti-ROS agent (Balasundaram et al., 1993; Chattopadhyay et al., 2003; Chattopadhyay et al., 2006; Ha et al., 1998a; Ha, et al., 1998b; Khan et al., 1992a; Khan et al., 1992b; Pegg, 2018; Stewart et al., 2018), a systematic dissection on the levels of individual ROS species in spermidine-enriched and spermidine-deficient conditions is missing. To address this, we incubated *E. coli* strains with 2’,7’-dichlorodihydrofluorescein diacetate (H2DCFDA), and dihydroethidium (DHE) probes, which generate fluorescent compounds reacting with one-electron-oxidizing species such as •OH, and O_2_**^-^** radicals, respectively (Chen et al., 2013; Kalyanaraman et al., 2012). The relative mean fluorescence intensity (MFI) of H2DCFDA was increased about 1.8-fold in the spermidine synthase-defective (Δ*speE*) strain, while significantly decreased in Δ*speG* strain (Figure 1A). Spermidine treatment further decreased the H2DCFDA fluorescence in Δ*speG* strain (Figure 1A). Similarly, the relative MFI of DHE probe was increased significantly (1.9-fold) in Δ*speE* strain of *E. coli* (Figure 1B). These findings are consistent with the observations that spermidine is an anti-ROS agent (Balasundaram et al., 1993; Chattopadhyay et al., 2003; Chattopadhyay et al., 2006; Ha et al., 1998a; Ha et al., 1998b; Khan, et al., 1992a; Khan et al., 1992b; Pegg, 2018; Stewart et al., 2018).

On the other hand, the relative MFI of DHE probe was increased significantly (2.3-fold) in the spermidine-fed Δ*speG* as compared to WT strain of *E. coli* (Figure 1B). Tiron (Tr), an O_2_**^-^** quencher, decreased the MFI of DHE in the spermidine-fed Δ*speG* strain (Figure 1B). These two observations, which indicate that the spermidine accumulation in the Δ*speG* strain evokes O_2_^-^ production, are in contrary to the generic idea that suggests spermidine is exclusively an anti-ROS agent (Balasundaram et al., 1993; Chattopadhyay et al., 2003; Chattopadhyay et al., 2006; Ha et al., 1998a; Ha et al., 1998b; Khan et al., 1992a; Khan et al., 1992b; Pegg, 2018; Stewart et al., 2018). In another assay, we determined that Δ*speE* and the spermidine-fed Δ*speG* strains release substantially low levels of H_2_O_2_ compared to the untreated counterpart and WT cells (Figure 1C).

Next, we allowed WT and Δ*speG* strains to grow against the spermidine-diffusing wells on agar plates in aerobic and anaerobic conditions (Figure 1D). A far wider zone of inhibition (ZOI) of growth for Δ*speG* strain was observed compared to WT under aerobic condition (Figure 1D), while a narrow ZOI was observed under anaerobic conditions for both strains (Figure 1D). This data confirms that O_2_^-^ production is the major cause of the observed spermidine toxicity.

If spermidine induces O_2_^-^ production, superoxide dismutase (SOD) genes (e.g., *sodA* and *sodB*) would play vital roles. Therefore, the serial dilutions of Δ*speG*Δ*sodA*, Δ*speG*Δ*sodB*, and Δ*speG*Δ*sodA*Δ*sodB* mutants were spotted on LB-agar plates to observe that all the double and triple mutants showed higher growth defects than Δ*speG* strain under spermidine stress (Figure 1E). However, the cell viability of the double mutants was similar to the Δ*speG* strain, while the triple mutant exhibited an accelerated loss of cell viability, in the presence of spermidine (Figure 1F). When we overexpressed *sodA* in the Δ*speG*Δ*sodA* strain from a plasmid, pBAD-*sodA*, the growth defect was suppressed (Figure 1E) and the cell viability was remarkably restored under spermidine stress (Figure 1F). Note that, unlike Δ*speG* strain, the single mutants show growth and viability similar to the WT strain in the presence or absence of spermidine (Figure 1E-F - figure supplement 1). This data confirms that the absence of SOD enzymes aggravates O_2_^-^ toxicity in the spermidine-fed Δ*speG* strain.

### SpeG-negative *S. aureus* also generates O_2_^-^ under spermidine stress

As mentioned earlier, *S. aureus* strains do not synthesize polyamines and usually they are *speG*-negative (Joshi, 2012). One such *speG*-negative *S. aureus*, RN4220 strain, encounters host-derived spermidine/spermine while invading host tissues. Unlike the RN4220 strain, the USA300 strain of *S. aureus* is a superior pathogen partly due to horizontal acquisition of *speG* in evolution (Joshi, 2012). Therefore, spermidine stress would likely evoke O_2_^-^ radicals in the *S. aureus* RN4220 but not in USA300 strain. Applying 3.2 mM exogenous spermidine, we confirmed that similar to the *E. coli* Δ*speG* strain, DHE fluorescence was significantly increased in *S. aureus* RN4220 strain, while spermidine treatment failed to elevate DHE fluorescence in the USA300 strain (Figure 2A). Furthermore, an extremely wide ZOI of growth of RN4220 as compared to the USA300 strain under aerobic condition indicate that spermidine is toxic in the absence of *speG* due to production of O_2_^-^ anions (Figure 2B). These results generalize our observation that spermidine-induced O_2_^-^ production exerts toxicity in bacteria.

**Figure 2.**
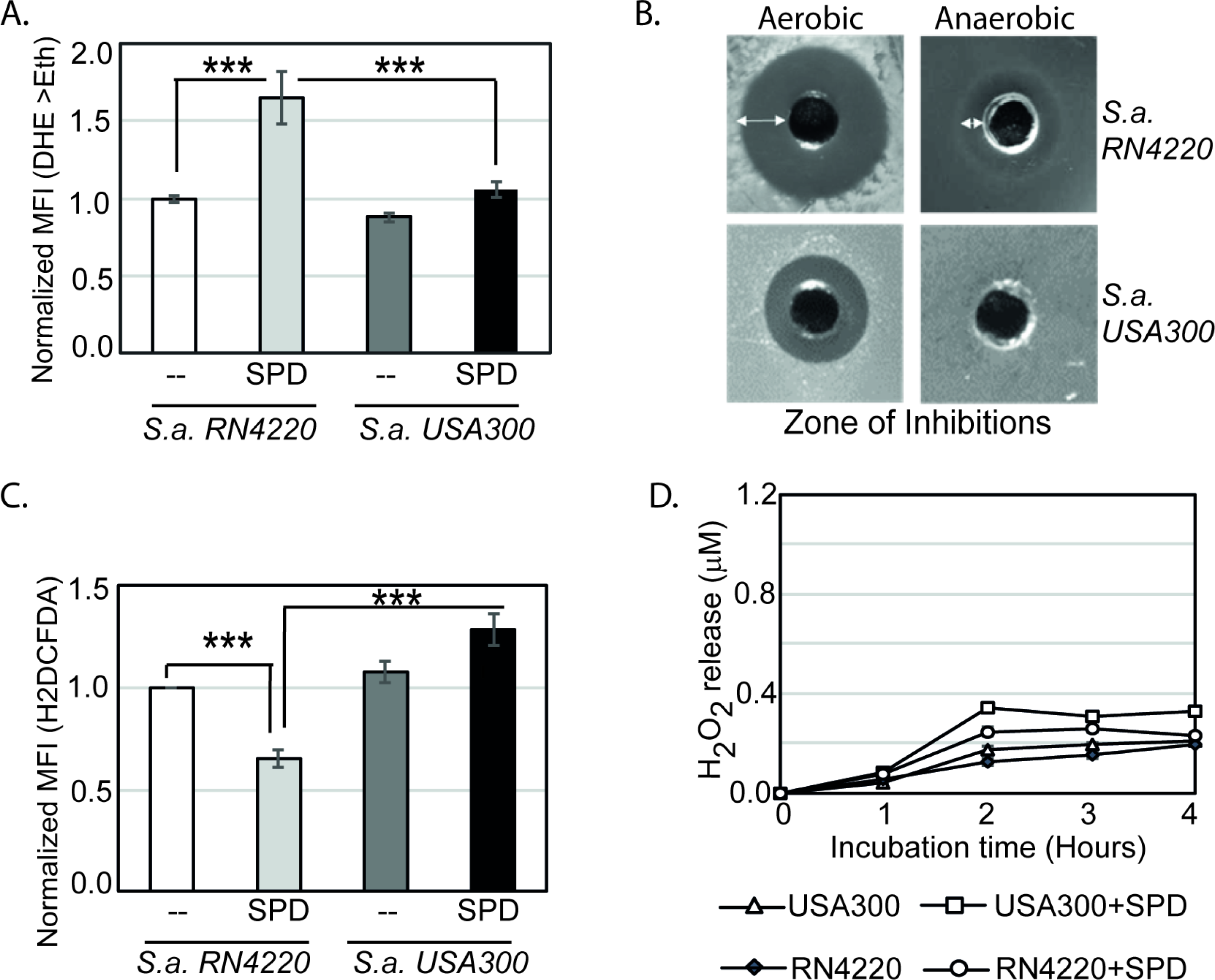
O_2_^-^ and other intracellular ROS production in *S. aureus*. **A.** The bar diagram shows normalized MFI of DHE fluorescence, as a function of O_2_**^-^** radical levels in *S. aureus* RN4220 and USA300 in the presence or absence of SPD. **B.** ZOI surrounding SPD well on the agar plate was determined for the *S. aureus* (*S.a.*) and USA300 strains under aerobic and anaerobic conditions. **C.** The bar diagram shows normalized MFI of H2DCFDA fluorescence DHE fluorescence, as a function of •OH radical levels in *S. aureus* RN4220 and USA300 in the presence or absence of SPD. **D.** The plot shows H_2_O_2_ production from *S. aureus* strains in the presence or absence of SPD for a span of 5 hours were shown. Error bars in the panels are mean ± SD from the three independent experiments. ** and *** denote P values <0.01 <001; unpaired T test. See also source data 5 and 6. **Source data 5-** Figure 2A and 2C raw data **Source data 6-** Figure 2D raw data

Similar to the results for *E. coli* Δ*speG* strain, spermidine also decreases the level of •OH radicals in RN4220 strain, as indicated by the modest decrease (0.65-fold) in H2DCFDA fluorescence (Figure 2C). However, no significant change in the H2DCFDA fluorescence was observed in USA300 strain (Figure 2C). In contrast to *E. coli* Δ*speG* strain, the *S. aureus* RN4220 and USA300 strains inherently produces low levels of H_2_O_2_ (Figure 2D; compare it with Figure 1C). Further spermidine treatment had no discernible effect on the H_2_O_2_ levels (Figure 2D).

### O_2_^-^ production under spermidine stress affects cellular redox state

Antioxidant chemicals, viz. Tiron (Tr), sodium pyruvate (SP), and thiourea (TU) scavenge O_2_^-^, H_2_O_2_, and •OH, respectively (Bleeke et al., 2004; Franco et al., 2007). Whereas, N-acetylcysteine (NAC) and ascorbate counterbalance oxidative stress replenishing glutathione levels and donating electrons to reducing partners (Nimse and Pal, 2015; Sun, 2010). We show that Tr, NAC, and ascorbate, but not SP and TU, rescued the spermidine-mediated growth inhibition phenotype (Figure 3A). This observation confirms that the O_2_^-^ stress perturbing redox environment is the route of spermidine toxicity.

**Figure 3.**
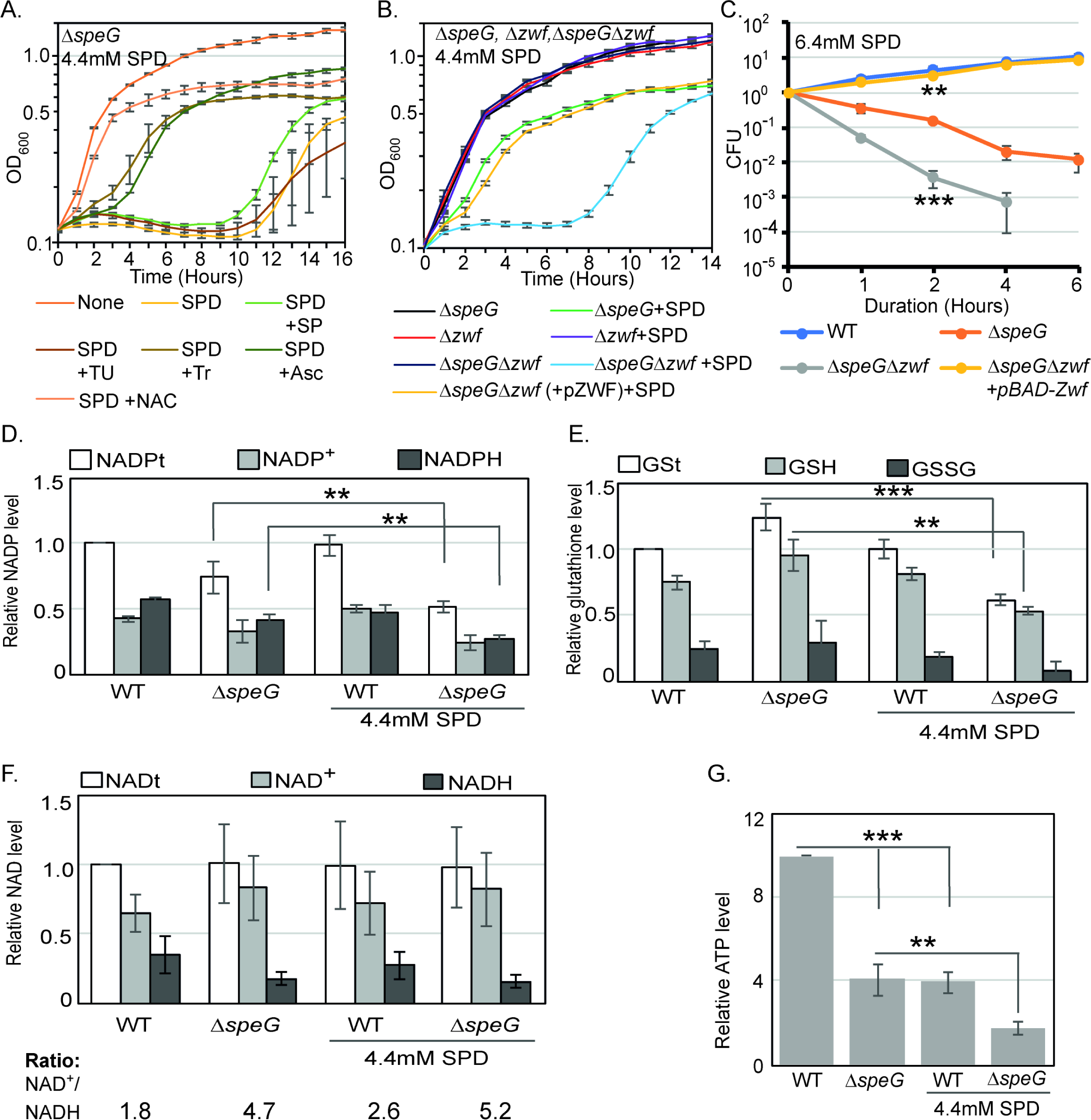
O_2_^-^ radical production affects redox balance in the spermidine-fed Δ*speG* strain. **A.** Growth curves show that Tiron (Tr), Ascorbate (Asc) and N-acetyl cysteine (NAC) can overcome spermidine (SPD) stress while sodium pyruvate (SP) and Thiourea (TU) fails to do so. Growth curves show that Δ*speG*Δ*zwf* strain is hypersensitive to SPD in comparison to Δ*speG* strain. Complementation of Δ*speG*Δ*zwf* strain with pZwf plasmid overcomes this SPD hypersensitivity. **C.** CFUs were obtained for different *E. coli* strains pretreated with SPD for desired time points and plotted to show the reduced viability of Δ*speG*Δ*zwf* strain in comparison to the Δ*speG* strain. **D.** Relative levels of NADPt and NADPH were significantly decreased in the Δ*speG* strain under SPD stress. **E.** Relative levels of GSt, GSH and GSSG were significantly decreased in the SPD-fed Δ*speG* strain. **F.** No significant change in the relative total NAD (NADt), NAD+ and NADH levels were recorded. However, NAD+ to NADH ratio was significantly increased in the Δ*speG* strain compared to WT cells. No further increase of the ratio was observed by adding SPD in the growth medium of WT and Δ*speG* strain. **G.** The relative level of ATP was declined in Δ*speG* strain and spermidine-fed WT cells in comparison to the unfed WT. SPD supplementation decreased the ATP level further in the SPD-fed Δ*speG* strain. Error bars in the panels are mean ± SD from the three independent experiments. Whenever mentioned, the *** and ** denote P values <0.001 and <0.01, respectively; unpaired T test. See also source data 7-11. **Source data 7-** Figure 3A raw data **Source data 8-** Figure 3B raw data **Source data 9-** Figure 3C raw data **Source data 10-** Figure 3D, 3E and 3F raw data **Source data 11-** Figure 3G raw data

The reduced nicotinamide adenine dinucleotide phosphate (NADPH) is a potent reducing agent. NADPH drives glutathione and thioredoxin cycles, thereby producing reduced forms of glutathione (GST), glutaredoxins, and thioredoxins to cope up with oxidative stress. A large fraction of NADPH in *E. coli* is provided by a Glucose-6-phosphate 1-dehydrogenase (Zwf) catalyzed reaction (Olavarría et al., 2012). We show that both the growth and viability of Δ*speG*Δ*zwf* double mutant were significantly affected compared to the Δ*speG* strain under spermidine stress (Figure 3B and 3C). Complementing Δ*speG*Δ*zwf* with a plasmid, pBAD-*zwf*, rescues the growth defect and mortality under spermidine stress (Figure 3B and 3C). We estimated the total NADP (NADPt), total glutathione (GSt), and their oxidized (NADP+ and GSSG) and reduced (NADPH and GSH) species in the WT and Δ*speG* strains grown in the absence and presence of spermidine. The relative levels of total and reduced species of NADP and GST were decreased significantly in the spermidine-fed Δ*speG* strain (Figure 3D and 3E). NAD serves as the precursor for NADP production. However, the levels of total (NADt), oxidized (NAD+), and reduced (NADH) did not alter significantly (Figure 3F). Nevertheless, the NAD+ to NADH ratio was significantly increased in the Δ*speG* strain compared to WT cells (Figure 3F). No significant increase of the ratios was observed by adding spermidine in the growth medium of WT and Δ*speG* strain (Figure 3F). In consistence with the increased ratio of NAD+ to NADH, the level of ATP was declined in Δ*speG* strain compared to the unfed WT (Figure 3G).

### Spermidine affects iron-sulfur cluster biogenesis and blocks the induction of SoxR regulon

To understand the global impact of spermidine toxicity, we performed a microarray experiment on the Δ*speG* strain in the presence and absence of spermidine. The genes that were >2-fold downregulated are involved in flagellar biogenesis, acid resistance, hydrogenase function, nitrogen metabolism, electron transport, aromatic and basic amino acid metabolism, etc. (Figure 4A and Supplementary file 1). Interestingly, transcription of the genes encoding chaperones, heat shock, and other stress factors (*groL*, *groS*, *dnaK*, *hdeAB*, *ibpAB*, *uspAB*, etc.) were also downregulated under spermidine stress (Supplementary file 1). On the other hand, among the highly upregulated category, the genes that encode for the ribosome, RNA polymerase, transcription factors, DNA polymerase, and enzymes for the fatty acid biosynthesis and iron-sulfur cluster (*isc*) biogenesis were prominent (Supplementary file 1 and Figure 4A). Many genes regulated by Fis and IHF were activated or repressed in our microarray indicating that spermidine could activate Fis and IHF regulon (Supplementary file 2). However, Δ*speG*Δ*fis*, but not Δ*speG*Δ*ihfA* strain, generated small colonies upon overnight incubation (Figure 4 – figure supplement 1), suggesting that the role of Fis regulator is critical under spermidine stress. RT-qPCR experiment was performed to validate the microarray data partially (Figure 4 – figure supplement 2).

**Figure 4.**
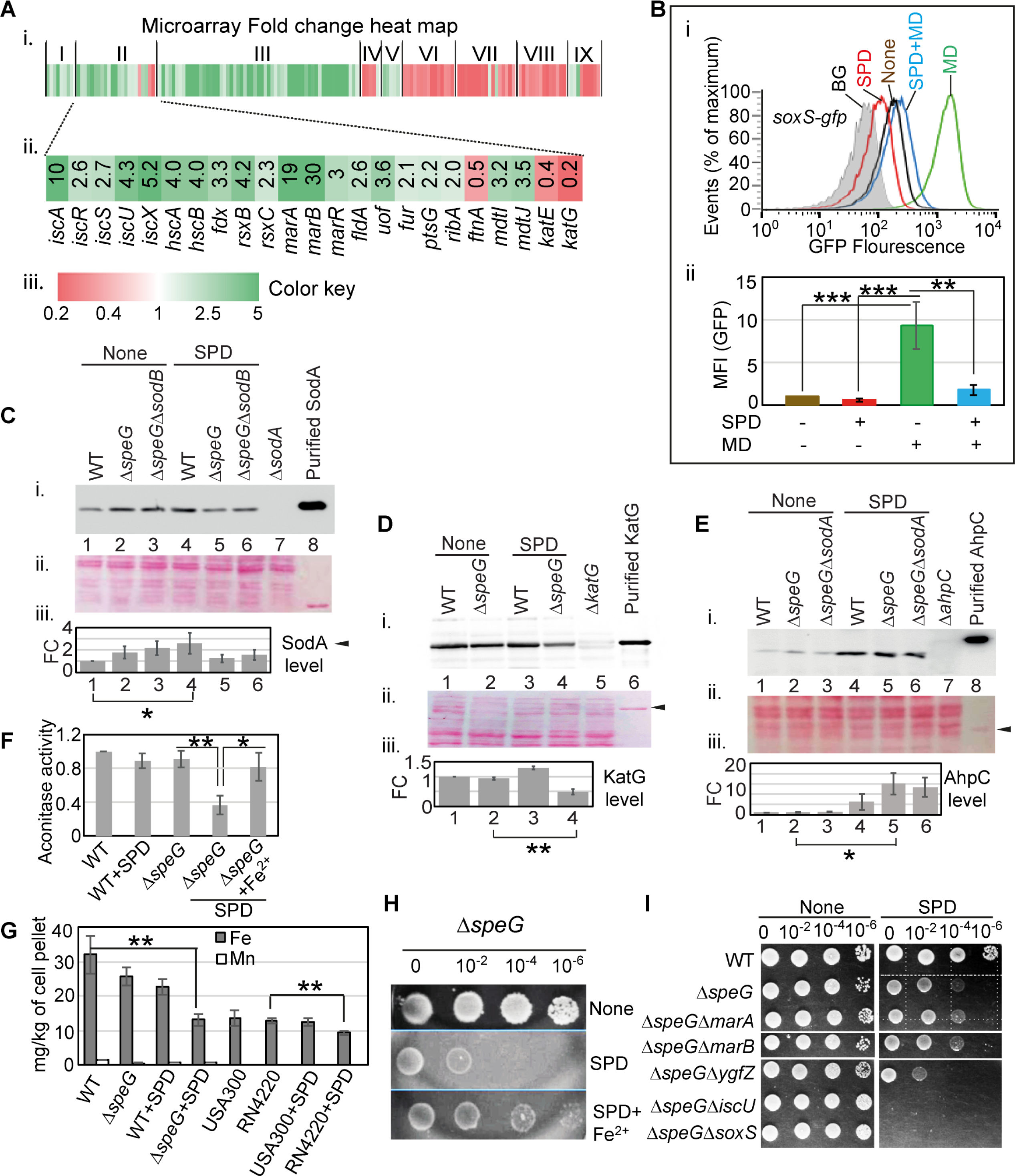
Spermidine blocks the activation of superoxide-defense circuit and affects iron metabolism. **A.** (i) Microarray heat map showing various categories of genes (categories: I-IX; see Supplementary file 1) that were differentially expressed under spermidine stress. (ii) Zoomed in heat-map of the category II genes responsible for iron metabolism and ROS regulation (iii) Color key represents the expression fold-change of the genes. **B.** The subpanel (i) represents a flow cytometry experiment to demonstrate that spermidine (SPD) stress inhibits menadione (MD)- induced P*soxS*-*gfpmut2* reporter fluorescence. The subpanel (ii) represents relative MFIs in the presence or absence of SPD and MD calculated from three different flow cytometry experiments. **C, D, E.** Western blotting experiments show SodA, KatG and AhpC levels in the various strains in the presence or absence of SPD; **(i)** developed blot, **(ii)** ponceau S-stained counterpart of the same blot, **(iii)** The bar diagrams represent relative fold change (FC) of the proteins under SPD stress. The relative FC values were calculated from the band intensity values obtained from three independent blots in comparison to the untreated WT counterparts. Purified 6X his-tagged SodA, KatG and AhpC proteins were loaded as positive controls. The cellular protein extracts from Δ*sodA*, Δ*katG* and Δ*ahpC* strains were used for negative controls. **F.** The bar diagram represents relative aconitase activity in the *E. coli* WT and Δ*speG* strains in the presence and absence of SPD. **G.** Intracellular levels of Fe and Mn levels for *E. coli* strains, and Fe levels for *S. aureus* strains determined in the presence or absence of SPD stress were plotted. **H.** Spot assay using serially diluted Δ*speG* cells demonstrated that Fe^2+^ can rescue SPD stress. **I.** Spot assay shows the relative sensitivity of various double mutants, Δ*speG*Δ*ygfZ,* Δ*speG*Δ*iscU* and Δ*speG*Δ*soxS* strains to SPD. Error bars in the panels are mean ± SD from the three independent experiments. Whenever mentioned, the ***, ** and * denote P values <0.001, <0.01 and <0.1 respectively; unpaired T test. See also Figure 4 – figure supplement 1, Figure 4 – figure supplement 2, and source data 12-16.

Iron-sulfur center of SoxR senses the levels of cellular O_2_^-^ or NO (Fujikawa et al., 2017; Hidalgo and Demple, 1994; Kobayashi, 2017; Liochev and Fridovich, 2011; Lo et al., 2012) and triggers transcription of a set of genes, including *soxS*, s*odA,* and *zwf* (Touati, 2000; Wu and Weiss, 1992). Surprisingly, none of the three critical genes was found to be activated in the microarray. RT-qPCR analyses verified the unaltered expression of *soxS*, *sodA,* and *zwf* under spermidine stress (Figure 4 – figure supplement 2). Consistently, using Δ*speG* harboring pUA66_*soxS*, a reporter plasmid expressing *gfpmut2* from the *soxS* promoter (P*soxS*-*gfpmut2*), and RKM1 strain containing a chromosomally fused *lacZ* reporter under *sodA* promoter (P*sodA*-*lacZ*) (Table 1), we did not find any transcriptional activation of *soxS* and *sodA* promoters (Figure 4 – figure supplement 2). Therefore, we suspected whether spermidine in excess blocks the O_2_^-^ -mediated activation of SoxR, thereby aggravating O_2_^-^ toxicity. To probe this possibility, we induced O_2_^-^ production by menadione to observe P*soxS*-*gfpmut2* reporter induction and chased it by spermidine in the Δ*speG* strain. Spermidine suppressed the menadione-induced GFP reporter fluorescence, suggesting that spermidine indeed blocks SoxR-mediated activation of *soxS* in *E. coli* (Figure 4B). A possible mechanism of spermidine-mediated SoxR inactivation is discussed. Among other ROS-responsive genes, the catalase coding genes (*katE* and *katG*) were downregulated, (Figure 4A) while no change was observed in the expression of *ahpCF* genes under spermidine stress (GEO accession #154618). Using pUA66_*ahpC* and pUA66_*katG* reporter plasmids (P*ahpC*-*gfpmut2* and P_*katG*_-*gfpmut2*, respectively), we validated these microarray observations (Figure 4 – figure supplement 2).

**Table 1.**
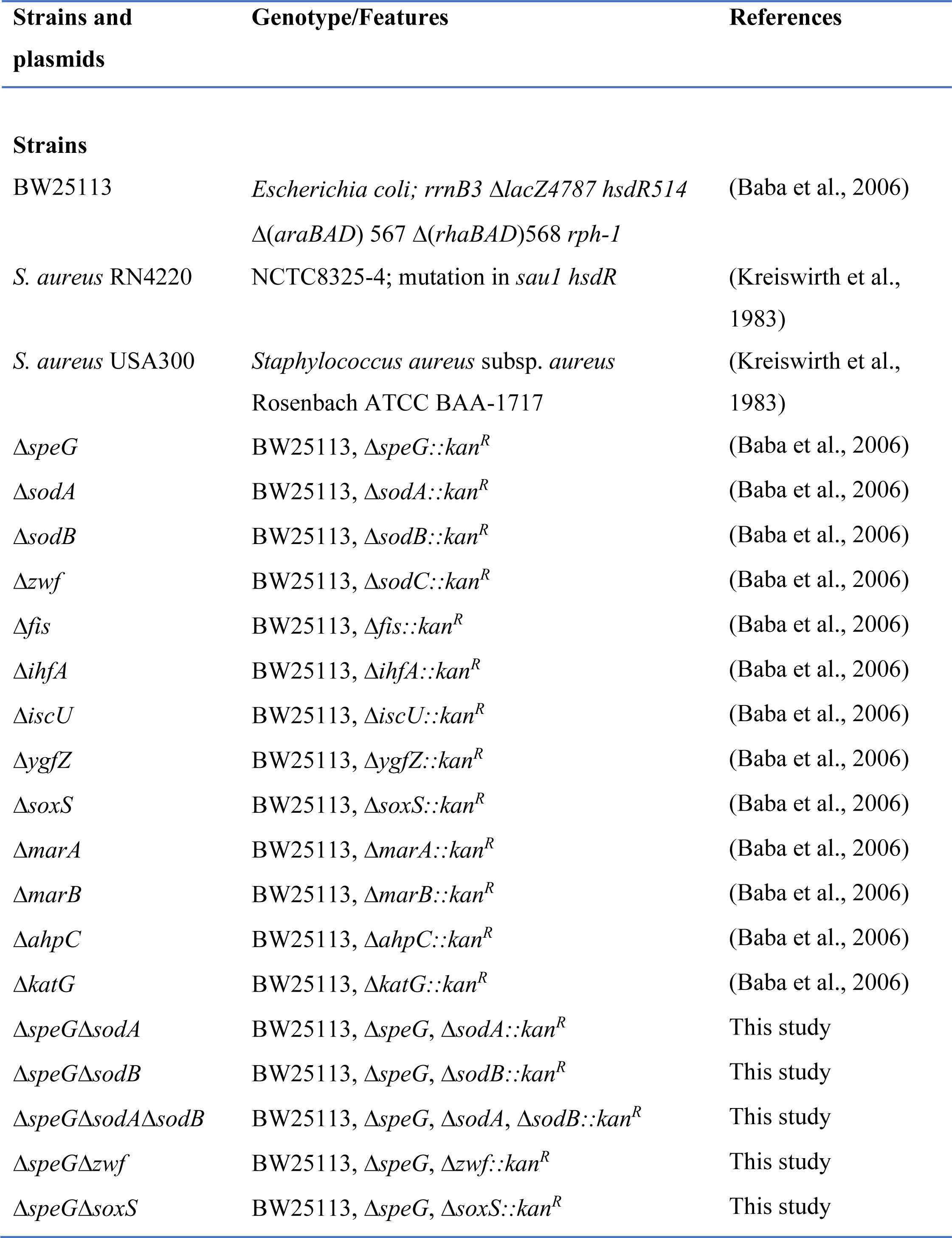
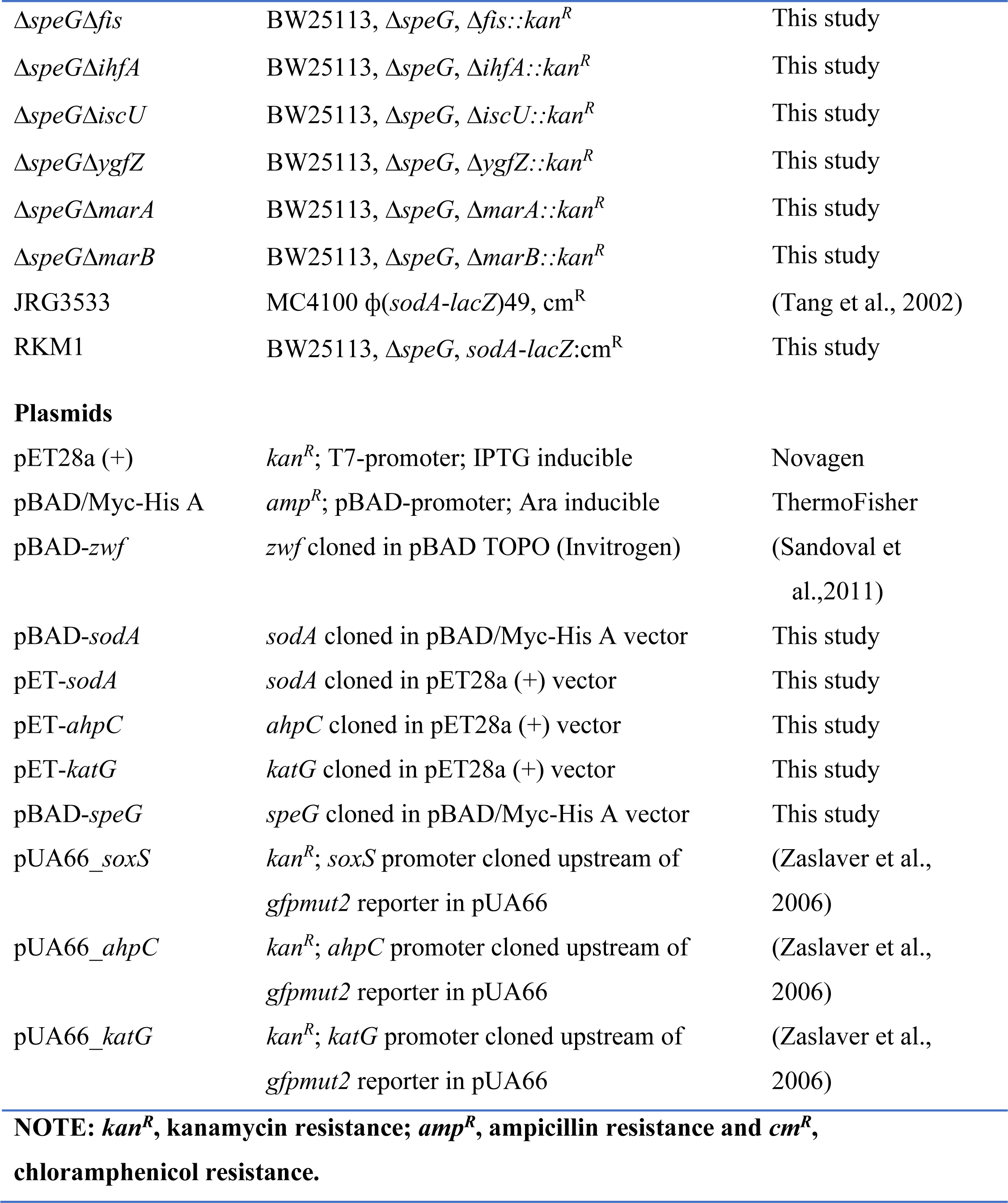
The list of strains and plasmids used in this work.

Consistent with the microarray expressions, our western blotting experiments exhibited the unchanged expression of SodA and a decreased expression of KatG in the spermidine-treated Δ*speG* strain compared to untreated counterparts (Figure 4C and 4D). However, SodA level was modestly elevated in the Δ*speG* strain, and the spermidine-treated WT strain, in contrast to the untreated WT strain (Figure 4C). Contrary to the microarray data, a profound increase in AhpC level was observed while growing WT or Δ*speG* cells in the presence of spermidine, indicating a translational elevation of AhpC level under spermidine stress (Figure 4E). Increased AhpC level indicating the activation of alkyl hydroperoxidase (AhpCF) enzyme, could be responsible for the decline in cellular H_2_O_2_ level (Figure 1C). Thus, declined H_2_O_2_ concentration could be the limiting factor for the cellular •OH radical production under spermidine stress (Figure 1A).

O_2_^-^ has the potential to oxidize the solvent-exposed iron-sulfur clusters of *E. coli* dehydratases, aconitase, and fumarase enzymes to liberate free Fe^2+^ (Benov, 2001; Fridovich, 1986; Imlay, 2008). Therefore, supplementation of Fe^2+^ ions helps to repair the damaged clusters (Gardner and Fridovich, 1992; Imlay, 2008). Consistently, we observed that the declined aconitase activity in the spermidine-stressed Δ*speG* strain was rescued by supplemental Fe^2+^ ion (Figure 4F). Besides, we show that the intracellular level of iron in the Δ*speG* strain was decreased more than three-fold in the presence of spermidine (Figure 4G). Interestingly, the iron content in the *S. aureus* strains was found to be substantially lower than the *E. coli* WT cells (Figure 4G). Furthermore, while the iron content of USA300 strain remained indifferent, the iron content of RN4220 strain was significantly declined under spermidine stress (Figure 4G). The iron scarcity was also reflected in the gene expression pattern of IscR regulon. IscR forms a functional holoenzyme with the iron-sulfur cluster. The de-repression of iron-sulfur cluster biogenesis operon (*iscRSUA-hscBA-fdx-iscX*) in the microarray (Figure 4A) signifies the presence of non-functional apo-IscR under the scarcity of cellular Fe^2+^ ion (Schwartz et al., 2001). Supplementation of Fe^2+^ salt in the LB-agar plate to rescue the growth of spermidine-fed Δ*speG* strain supports this claim (Figure 4H). Spermidine also activated *rsxA* and *rsxB* (Figure 4A), which encode the critical components of the iron-sulfur cluster reducing system of SoxR (Koo et al., 2003). The level of manganese, an antioxidant metal that determines *sodA* activity, is usually increased under iron scarcity (Kaur et al., 2014; Kaur et al., 2017; Martin et al., 2015; Waters et al., 2011). However, a modest decrease in the level of cellular manganese under spermidine stress was observed (Figure 4G). The low level of manganese could slow down the rate of dismutation of O_2_^-^ anion compromising SodA function, thereby elevating the O_2_^-^ anion levels in the spermidine-treated cells. Finally, we spotted the cultures of serially diluted *E. coli* strains to show that the deletion of two individual genes (*iscU* and *ygfZ*), which are involved in the iron-sulfur cluster biogenesis (Waller et al., 2010), affects the growth of the spermidine-treated Δ*speG* strain (Figure 4I). Interestingly, the Δ*speG*Δ*soxS* strain was more sensitive to spermidine than the Δ*speG* strain (Figure 4I), indicating that the basal level of *soxS* expression has some potential to ameliorate O_2_^-^ under spermidine stress. Although *marA* and *marB* genes were expressed at the highest level in the spermidine-stressed Δ*speG* strain (Figure 4A), Δ*speG*Δ*marA* and Δ*speG*Δ*marB* strains did not show any difference in growth compared to Δ*speG* strain under spermidine stress (Figure 4I). Note that, unlike Δ*speG* strain, the single mutants grow similarly to the WT strain in the presence or absence of spermidine (Figure 4I and Figure 1-figure supplement 1).

### Free spermidine interacts and oxidizes Fe^2+^ ion to generate superoxide radicals in vitro

To probe whether spermidine directly interacts with iron, we performed isothermal titration calorimetry (ITC) using Fe^3+^ (Ferric citrate) and Fe^2+^ (Ferrous ammonium sulfate) ions. Titration of spermidine with Fe^3+^ generated exothermic peaks indicating a standard binding reaction with a stoichiometry (N) of 0.711 (Figure 5A). On the other hand, titration of spermidine with Fe^2+^ in two different isothermal conditions produced consistent and complex patterns (Figure 5B and 5C). To explain it, we divided the pattern into two halves. In the first half, Fe^2+^ injections to spermidine generated alternate exothermic and endothermic peaks till the ratio of spermidine to Fe^2+^ reaches about 1:1.3 (Figure 5B and 5C). In the second half of the profile, after the ratio of spermidine to Fe^2+^ crosses 1:1.3, no endothermic peaks were observed, and a gradual shortening of exothermic peaks was generated, leading to saturation (Figure 5B and 5C). From the first half of pattern, we suspected Fe^2+^ interaction with spermidine also involves some other reactions, such as oxidation of the Fe^2+^ to generate Fe^3+^ and O_2_^-^, Fe^3+^ release, and subsequent Fe^3+^ binding to spermidine.

**Figure 5.**
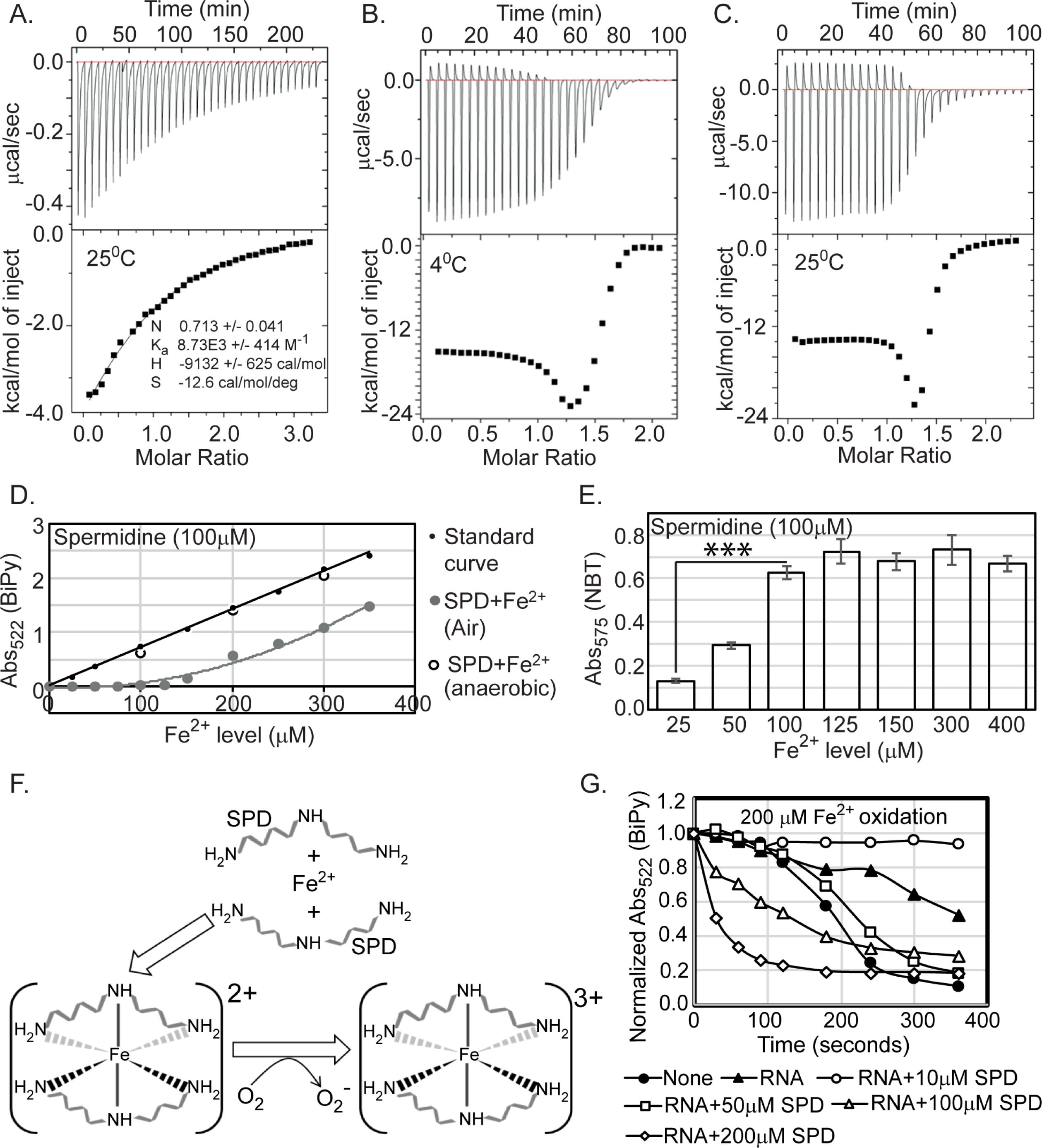
Spermidine oxidizes Fe^2+^ generating O_2_^-^ radical in aerobic condition. **A.** ITC data demonstrates the interaction of spermidine with Fe^3+^. **B** and **C.** ITC data shows the interaction of spermidine with Fe^2+^ ion at 4°C and 25°C, respectively. **D.** 100 µM spermidine was incubated with different concentrations of Fe^2+^ followed by estimation of Fe^2+^ levels by bipyridyl chelator. The color formation was recorded at 522 nm and plotted them along with standard curve. The panel depicts that the incubations of 100 µM spermidine with 100, 200 and 300 µM of Fe^2+^ in the anaerobic condition do not lead to the loss of Fe^2+^ ions detected by bipyridyl chelator. However, when 100 µM spermidine was incubated with the different concentrations of Fe^2+^ (25 µM to 350 µM) in the aerobic condition, the bipyridyl-mediated color formation was observed when Fe^2+^ level was between above 125 µM to 150 µM (i.e., till spermidine to Fe^2+^ ratio reaches approximately 1.3). The mean values from the three independent experiments were plotted. SD is negligible and is not shown for clarity. **E.** NBT assay was performed to determine that spermidine and Fe^2+^ interaction yields O_2_^-^ radical. The colorimetry at 575 nm suggests that 100 µM of spermidine interacts approximately with 125 µM of Fe^2+^ (ratio 1:1.3) to generate saturated color. Error bars in the panel are mean ± SD from the three independent experiments. *** denotes P value <0.001; unpaired T test. **F.** Model to show final coordination complex formation. An Fe^2+^ interacts with two spermidine molecules forming hexadentate co-ordination complex. This interaction oxidizes Fe^2+^ liberating one electron to reduce oxygen molecule. Finally, two spermidine coordinates one Fe^3+^ with an octahedral geometry. **G.** The curves represent the *E. coli* total RNA inhibits iron oxidation. Spermidine further reduces the RNA-mediated iron oxidation at concentration 10 µM but higher concentrations of spermidine increase the iron oxidation despite the presence of RNA. The mean values are derived from the three independent experiments and plotted. SD is negligible and is not shown for clarity. See also source data 17-19. **Source data 17-** Figure 5D raw data **Source data 18-** Figure 5E raw data **Source data 19-** Figure 5G raw data

To test whether Fe^2+^ was oxidized in the presence of spermidine to liberate Fe^3+^, we titrated spermidine by increasing amounts of Fe^2+^ iron followed by assessing the level of Fe^2+^ by using bipyridyl chelator. Chelation of Fe^2+^ ions by bipyridyl generates pink color indicating Fe^2+^ levels. No color formation was observed till the ratio of spermidine to Fe^2+^ reaches 1:1.3 (Figure 5D), a number that exactly matches with the ratio of spermidine to Fe^2+^ in the first half of ITC experiments (Figure 5B and 5C). The color formation starts appearing when the ratio crosses 1:1.3 (Figure 5D), suggesting that 1 molecule of spermidine (or 10 molecules) exactly oxidizes 1.3 molecules (or 13 molecules) of Fe^2+^. The colorimetric values overlap with the standard curve when reactions were under anoxic condition, indicating Fe^2+^ was not oxidized (Figure 5D). We used nitro blue tetrazolium (NBT) dye to check whether the loss of one electron from Fe^2+^ generates O_2_^-^ anion under spermidine stress. An increased NBT absorption at 575nm till the ratio of spermidine to Fe^2+^ reaches 1:1.3 confirms that 1 molecule (or 10 molecule) of spermidine interacts with 1.3 molecules (or 13 molecules) of Fe^2+^ generating 1.3 molecules (or 13 molecules) O_2_^-^ anion radical (Figure 5E). From the stoichiometry of 0.711 (which is close to 0.5) (Figure 5A), we postulate that two spermidine and one Fe^3+^ together could form a hexadentate co-ordination complex with an octahedral geometry (Figure 5F). It appears that when spermidine molecules engaged to form a hexadentate co-ordination complex with Fe^2+^, the former helps oxidizing latter to form Fe^3+^ in sufficient concentrations. Fe^3+^ finally forms coordination complex with spermidine (Figure 5F). It may be noted that the binding of spermidine and Fe^3+^ is entirely enthalpy driven, as indicated by a large negative ΔH. The negative entropy (ΔS) value presumably results from the ordering of spermidine from an extended conformation to a compact and rigid one after metal chelation (Figure 5A).

The cellular spermidine barely exists as a “free” species; rather, majority of them remain “bound” with RNA, DNA, nucleotides, and phospholipids (Igarashi and Kashiwagi, 2000; Miyamoto et al., 1993; Schuber, 1989; Tabor and Tabor, 1984). It has been reported that these phosphate-containing biomolecules have the inherent property to inhibit iron oxidation blocking O_2_^-^ production (LØVaas, 1996; Tadolini, 1988a; Tadolini, 1988b). The bound spermidine further enhances the inhibitory effects of these biomolecules towards iron oxidation. Consistent with the report (Tadolini, 1988b), we noticed that 1 μg of RNA inhibited the oxidation of 200 µM Fe^2+^. The presence of 10μM spermidine further decreased iron oxidation (Figure 5G). However, increasing the concentrations of spermidine (50, 100, and 200 μM), accelerated iron oxidation gradually (Figure 5G). This data clearly indicates that cell maintains a level of cellular spermidine that may remain optimally bound with the biomolecules inhibiting O_2_^-^ generation. However, when homeostasis fails due to *speG* deletion, excess spermidine accumulates that can remain in a “free”- form inducing O_2_^-^ radical toxicity.

## Discussion

Our study presented in this paper answers why spermidine homeostasis is intriguingly fine-tuned in bacteria. We provide clear-cut evidence that excess spermidine, which remains as a free species (Figure 5G), stimulates the production of toxic levels of O_2_^-^ radicals in *E. coli* and *S. aureus*, unless spermidine is inactivated by the SpeG-mediated acetylation (Figure 1 and Figure 2). O_2_^-^ anion thus generated affects cellular redox balance (Figure 3) and damages iron-sulfur clusters of the proteins (Figure 4). Since spermidine directly interacts with Fe^2+^ (Figure 5), it may abstract iron from some of the iron-sulfur clusters, thereby inactivating some of the proteins. On the other hand, when spermidine level is at optimum, most of it remain as bound form with the biomolecules, thereby slows down iron oxidation and subsequent O_2_^-^ production (LØVaas, 1996; Tadolini, 1988a; Tadolini, 1988b) (Figure 5G). Thus, spermidine-deficiency would enhance the rate of iron oxidation (Figure 5G), leading to the production of O_2_^-^ and •OH radicals (Figure 1A and 1B). This is why spermidine is a double-edged sword where in excess, it provokes O_2_^-^ anion production, and in scarcity, it leads to increased O_2_^-^ anion and •OH radical production.

Polyamines remain protonated at physiological pH, yet they are able to coordinate several positively charged metal ions, such as Ni^2+^, Co^2+^, Cu^2+^, and Zn^2+^, possibly via charge neutralization by counterions that reduces the Coulombic repulsion between spermidine and the metals (LØVaas, 1996). Similar charge neutralization of the nitrogen atoms of spermidine likely allows coordinate covalent bonds with Fe^3+^ (Figure 5F). About 10 spermidine molecules oxidize Fe^2+^ to generate 13 Fe^3+^ cations and equivalent numbers of O_2_^-^ radicals (Figure 5B and 5C). When sufficient concentration of Fe^3+^ is generated, two spermidine molecules coordinate one Fe^3+^ to form a hexadentate complex with an octahedral geometry (Figure 5F). We substantiated this in vitro spermidine-mediated iron oxidation and subsequent O_2_^-^ radical production phenomena (Figure 5), showing that cells are highly toxic to the spermidine under aerobic condition but not under anaerobic condition (Figure 1J and 2B).

Usually, abundant O_2_^-^ level leads to H_2_O_2_ and •OH production. However, despite elevated O_2_^-^ production, spermidine lowers H_2_O_2_ and •OH levels in Δ*speG* strain. (Figure 1C and 1A). The declined H_2_O_2_ level could be attributed to the slower rate of O_2_^-^ anion dismutation due to the failure of *sodA* activation (Figure 4C – figure supplement 2) and the activation of alkyl hydroperoxidase (AhpCF) that neutralizes H_2_O_2_, represented by AhpC overexpression (Figure 4D). A low level of cellular manganese (Figure 4G) could also limit SodA activity. Besides, the activation of IscR regulon (Figure 4A), the low cellular iron content (Figure 4G), and the rejuvenation of cell growth by Fe^2+^ supplementation (Figure 4H) indicate that the spermidine presumably lowers the Fe^2+^/Fe^3+^ ratio in Δ*speG* strain. Thus, the decreased level of Fe^2+^ and H_2_O_2_ (Figure 4G and 1C) could potentially diminish cellular •OH radical production in the spermidine-fed cells (Figure 1A). We have summarized all these observations and hypotheses in the schematic Figure 6.

**Figure 6.**
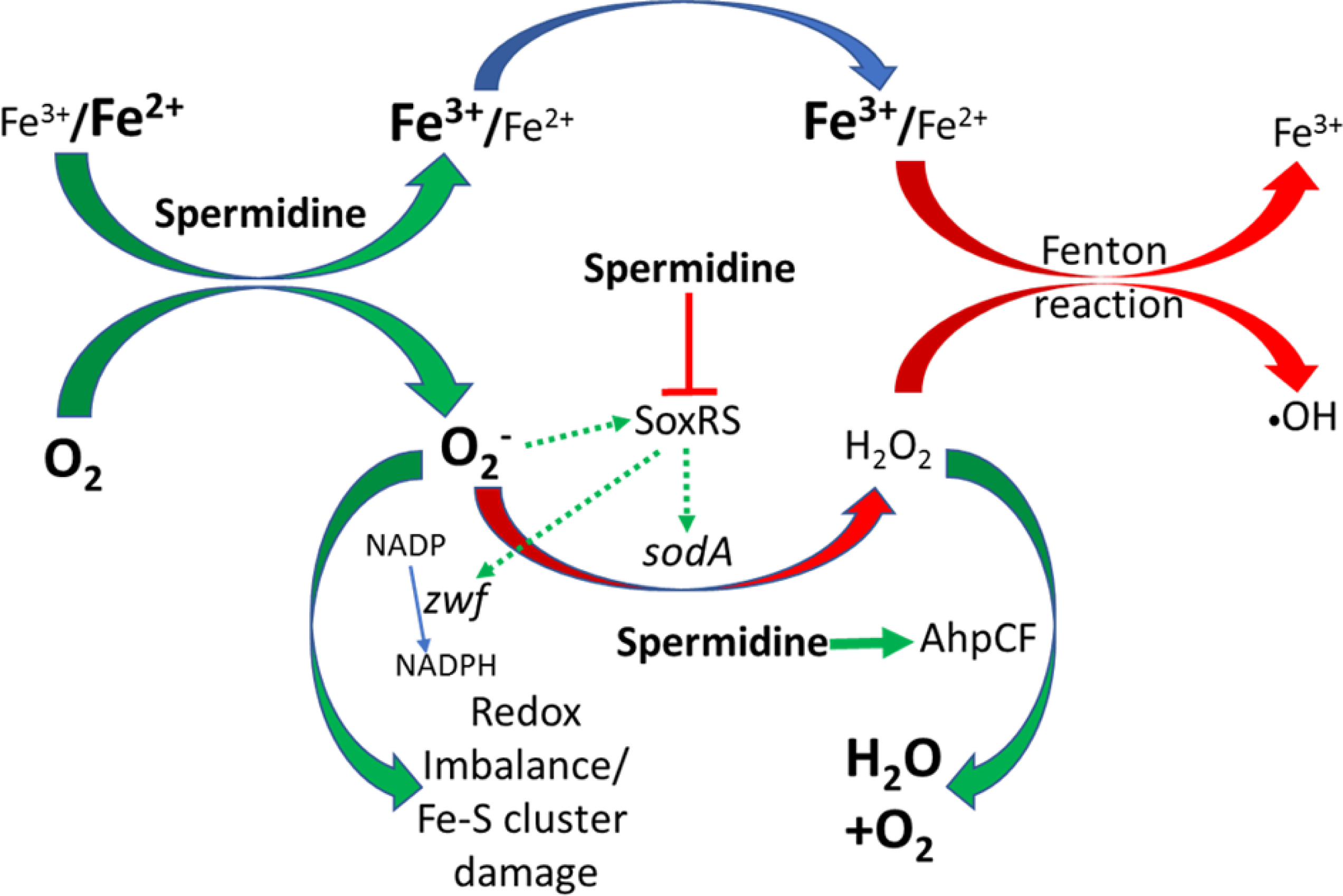
Flow-chart explaining the ROS generation under spermidine stress. The model describes that the spermidine administration in the cell interacts with free iron and oxygen to generate O_2_^-^ radical, increasing Fe^3+^/Fe^2+^ ratio. Spermidine also blocks O_2_^-^ radical-mediated activation of SoxRS that upregulates *zwf* and *sodA*. Consequently, NADPH production and dismutation of O_2_^-^ radical to H_2_O_2_ were not accelerated, leading to redox imbalance and O_2_^-^ - mediated damage to the iron-sulfur clusters, respectively. Additionally, spermidine translationally upregulated alkyl hydroperoxidase (AhpCF) that lowers the level of H_2_O_2_. Declined cellular Fe^2+^ and H_2_O_2_ levels weaken Fenton reaction to produce •OH radical. **Supplementary file 1.** Table listing microarray data representing upregulated (green) and downregulated (red) genes **Supplementary file 2.** Table listing the Fis and IHF regulated genes that were upregulated (green) and downregulation (red). **Supplementary file 3.** Table listing the oligonucleotide primers used in this study.

Interestingly, spermidine stimulates O_2_^-^ production and blocks O_2_^-^ -mediated SoxR activation in the Δ*speG* cells (Figure 4B). Since spermidine ubiquitously interacts with DNA and modulates gene expression in many ways (Igarashi and Kashiwagi, 2000; Jung and Kim, 2003; Miyamoto et al., 1993), one possibility could be that excess of it might occlude SoxR-binding to the *soxS* and *sodA* promoter regions to activate them. Alternatively, blockage of SoxR activation could result from spermidine-mediated activation of *rsxA* and *rsxB* (Figure 4A), which encode the critical components of the iron-sulfur cluster reducing system of SoxR (Koo et al., 2003), to keep SoxR inactive. The third possibility could be that the iron-binding property of spermidine (Figure 5A, 5B, 5C) may allow it to directly abstract iron from the iron-sulfur cluster to inactivate SoxR. Nevertheless, a detailed biochemical study on this aspect is needed to understand the mechanism. Our study clarifies how the horizontal acquisition of *speG* gene could confer a pathogenic advantage to the *S. aureus* USA 300 strain (Eisenberg et al., 2009). *S. aureus*, a Gram-positive commensal living on human skin, often causes severe disease upon access to deeper tissues. Since most of the iron in mammals exists intracellularly, the extracellular pathogen, *S. aureus* faces hardship and competes with the host for the available iron (Hammer and Skaar, 2011). As spermidine declines cellular iron content and interferes with iron metabolism (Figure 4), it is thus possible that *S. aureus* does not synthesize spermidine. Furthermore, the acquisition of *speG* gene could allow S. *aureus* USA300 to inactivate host-originated spermidine/spermine, thereby to maintain cellular iron content (Figure 4G). Corroborating to our findings, a recent observation has pointed out that spermine stress upregulates iron homeostasis genes, indicating that spermine toxicity has a specific connection with iron depletion in the *speG*-negative *S. aureus* strain, Mu50 (MRSA) (Yao and Lu, 2014). Besides, spermine-mediated iron depletion may be responsible for the synergistic effect of spermine with the antibiotics against *S. aureus* (Kwon and Lu, 2007). Nevertheless, a thorough in vivo host-pathogen interaction study will unravel a specific link between spermine/spermidine and iron depletion in *S. aureus*.

## Methods

### Bacterial strains, plasmids, proteins, and chemicals

Bacterial strains and plasmids used in this study are listed in Table 1. BW25113 strain of *E. coli* was used as WT in this study. *S. aureus* RN4220 strain is a gift from Ravi PN Mishra lab, CSIR IMTECH, India. *S. aureus* USA300 strain was purchased from ATCC (ATCC BAA-1717). Oligonucleotides were purchased from IDT. Bacterial broths and agar media were purchased from BD Difco. The knockout strains of *E. coli* were procured from the KEIO library (Baba et al., 2006), verified by PCR, and freshly transduced into the WT background by P1 phage. The double and triple knockout mutants were generated following the standard procedure described by Datsenko and Wanner (Datsenko and Wanner, 2000). *E. coli* strain JRG3533 was a generous gift from Dr. Rachna Chaba, IISER Mohali, India. RKM1 strain was constructed by P1 transduction of *sodA-lacZ*:Cm^R^ genotype of JRG3533 to BW25113Δ*soxS* strain.

The plasmids, pUA66_*soxS*, pUA66_*ahpC*, pUA66_*katG* were the gifts from Dr. Csaba Pal, Biological Research Centre of the Hungarian Academy of Sciences (Zaslaver et al., 2006). pBAD-*zwf* was a generous gift from Dr. CC. Vasquez, Universidad de Santiago de Chile (Sandoval et al., 2011). *sodA*, *katG*, *ahpC*, and *speG* genes were PCR amplified by DG12-DG13, RM7-RM8, DG9-DG10, and RK3-RK4 primer pairs (Supplementary file 3), respectively. The PCR products were double-digested at the primer-specific unique restriction sites and inserted into identically digested pET28a (+) plasmid vector so that the 6X his-tagged SodA, KatG, and AhpC proteins are being produced. The protein expression vectors, pET-*sodA*, pET-*ahpC*, pET-*katG* were transformed to BL21 (DE3) cells, and expressions were induced by 0.4 mM IPTG. The overexpressed proteins were purified using Ni-NTA beads. The purified proteins were used to raise rabbit polyclonal antibodies following the standard procedure. *sodA* and *speG* were additionally subcloned in pBAD/Myc-His A vector to get *sodA* and *speG* multicopy expressions for complementation assays.

### Growth, viability, spermidine sensitivity, and complementation assays

An automated BioscreenC growth analyzer (Oy growth curves Ab Ltd.) was used to generate growth curves mentioned in the results. For this purpose, overnight cultures of different strains were diluted in fresh LB medium and grown in the presence and absence of 3.2 mM to 6.4 mM of spermidine. 10 mM of each of the ROS quenchers (thiourea, Tiron, sodium pyruvate, ascorbate, and NAC) were used wherever mentioned. For viability assays, serially diluted *E. coli* strains were spread on LB-Agar surface supplemented with 6.4 mM spermidine. We determined the viability under spermidine stress from the number of colonies grown. Zone of inhibitions (ZOI), which appeared following overnight growth of the strains in the presence of 6.4 mM spermidine in the wells on agar plates, were determined both in aerobic and anaerobic conditions. The anaerobic condition was created in an anaerobic petri dish jar using AnaeroGas Pack 3.5L pouches. For complementation experiments, the pBAD-*zwf* and pBAD-*sodA* plasmids were transformed into Δ*speG*Δ*zwf,* and Δ*speG*Δ*sodA* strains, respectively, and growth assays were performed in the presence of spermidine. Since the leaky expressions of *zwf* and *sodA* were sufficient to rescue growth defects, induction with arabinose was avoided for this purpose.

The reporter plasmids, pUA66_*soxS*, pUA66_*ahpC*, pUA66_*katG*, were transformed in Δ*speG* strain. The transformed cells were grown in the presence or absence of 3.2 mM spermidine. Wherever mentioned, 25 µM menadione was used as a positive control for O_2_^-^ generation. The cell pellets were washed twice with PBS and dissolved in 500 µl phosphate buffer saline (PBS). Flow cytometry was done using the Fl1 laser for 0.05 million cells using FACSVerse (BD Biosciences). The mean fluorescence intensity (MFI) values from three biological replicates have been calculated.

### Determining relative ROS levels in the cells

H2DCFDA (10 µM) and DHE (2.5 µM), were used to measure cellular •OH and O_2_**^-^** anion, respectively. The cells were grown in the presence or absence of 3.2 mM spermidine. Cells were harvested, washed with PBS, and an equal mass of cell pellets was incubated with DHE or H2DCFDA probes for an hour. The data were acquired using BD accuri Fl3 laser (for DHE) and Fl1 laser (for H2DCFDA) for 0.05 million cells. The mean fluorescence intensities (MFI) values of triplicate experiments were calculated. For H_2_O_2_ detection, the *E. coli* cells were grown in the presence or absence of 3.2 mM of spermidine for 4 hours. Cells were harvested and washed with 1X M9 minimal media. The equal mass of cells (2.5 mg each) suspended in 6 ml M9 minimal media were incubated for different time points to allow H_2_O_2_ liberation. The relative H_2_O_2_ liberation was measured by a Fluorimetric Hydrogen Peroxide Assay kit (Sigma Aldrich).

### β-galactosidase and GFP reporter assays

For the β-galactosidase assay, the RKM1 strain was grown in the presence or absence of 3.2 mM of spermidine. The cell pellets were washed twice with Z-buffer (60 mM Na_2_HPO_4_, 40 mM NaH_2_PO_4_, 10 mM KCl, and 1 mM MgSO4) and diluted to OD600 ∼ 0.5. Promoter activity was measured by monitoring β-gal expression from single-copy *sodA-lacZ* transcriptional fusion. 100 µl of 4 mg/ml ONPG was used as a substrate, which was cleaved by β-galactosidase to produce yellow-colored O-nitrophenol. Colorimetric detection of this compound was done at 420nm.

The reporter plasmids, pUA66_*soxS*, pUA66_*ahpC*, pUA66_*katG*, containing GFP-mut2 reporters, were used to determine the promoter activities of *soxS*, *ahpC,* and *katG* genes in the presence or absence of 3.2 mM spermidine. Flow cytometry was done using the FL1 laser for 0.05 million cells using FACSVerse (BD Biosciences) or BD Accuri™ C6 Plus Flow Cytometer (BD Biosciences) machine.

### Western blotting experiments

Overnight culture of *E. coli* strains was inoculated in fresh LB medium in 1:100 dilution and grown for 1.5 hours at 37°C. Next, 3.2 mM of spermidine were added, wherever required and allowed to grow again at 37°C for 2.5 hours. Cells were harvested and lysed with B-PER^®^ bacterial protein extraction reagent (Thermo Scientific). The total protein level was checked by the Bradford assay kit (Bio-Rad). 40 µg of total cellular proteins from the individual samples were subjected to SDS-PAGE. The proteins were transferred to a nitrocellulose membrane and stained with Ponceau S to visualize protein resolution and equal loading in the PAGE. Western blotting was performed using polyclonal rabbit primary antibodies and HRP conjugated secondary antibodies. The blots were developed by Immobilon^®^ Forte Western HRP substrate (Millipore).

### Isothermal titration calorimetry

A MicroCal VP-ITC calorimeter, MicroCal Inc., was used for calorimetric measurements to probe the interaction of spermidine with Fe^2+^ and Fe^3+^ species. In order to achieve this, 100 µM of spermidine solution was prepared in 20 mM sodium acetate buffer (pH 5.5) and put into the sample cell. The ligands, 2.1 mM of FeCl3 or ferrous ammonium sulphate, were also dissolved in the identical sodium acetate buffer. The titrations involved 30 injections of individual ligands (5 µl per shot) at 300 seconds intervals into the sample cell containing 1.8 ml of 100 µM spermidine. The titration cell was kept at some specific temperature and stirred continuously at 286 rpm. The heat of dilution of ligand in the buffer alone was subtracted from the titration data. The data were analyzed using Origin 5.0 software.

### 2,2′-Bipyridyl and NBT assays

2,2′-Bipyridyl chelates Fe^2+^ producing color that absorbs at 522 nm (A522). The standard curve for 0 µM to 350 µM of Fe^2+^ ion was generated simply by recording A522 in the presence of 2,2′-Bipyridyl. Dissolved oxygen of medium and headspace oxygen was replaced by flushing N_2_ gas in the medium for 5 minutes to create an anoxic condition as described (Stieglmeier et al., 2009). To check whether spermidine acts as a catalyst for Fe^2+^ to Fe^3+^ oxidation, we performed 2, 2′-bipyridyl assay probing leftover Fe^2+^ after the reaction. For this assay, 100 µM of spermidine was incubated with increasing concentrations (25 µM to 350 µM) of ferrous ammonium sulphate for 10 minutes at room temperature. 900 µl of the reaction products were mixed with 90 µl 4M sodium acetate buffer (pH 4.75) and 90 µl bipyridyl (0.5% in 0.1N HCl). The color formation was recorded at 522 nm (A522) using UV-1800 Shimandzu UV-spectrophotometer. In another experiment, the assay was performed in anoxic condition using rubber capped sealed glass vials containing anoxic reactants and needle-syringe-mediated mixing of the reagents. Here, three different concentrations (100 µM, 200 µM, and 300 µM) of ferrous ammonium sulphate were reacted with 100 µM of spermidine for 10 minutes followed by spectrophotometry at A522. The standard curve for 0 µM to 350 µM of Fe^2+^ ion was generated simply by recording A522 of the mixture of 900 µl ferrous ammonium sulphate, 90 µl sodium acetate buffer, and 90 µl bipyridyl solutions.

Iron oxidation in the presence of RNA and spermidine was performed as described (Tadolini, 1988b). 1 µg RNA and increasing concentrations of spermidine (10 µM-200 µM) were used in 5 mM MOPS buffer, pH 7.4. The oxidation was started adding 200 µM FeCl2. The reactions were stopped at desired time point by adding a stop solution (1:1 4M sodium acetate:4M glacial acetic acid) followed by 2,2’-Bipyridyl to detect Fe^2+^ levels.

We used Nitro blue tetrazolium (NBT) dye to probe whether spermidine-stimulated Fe^2+^ to Fe^3+^ oxidation liberates O_2_^-^ anion *in vitro*. For this assay, different concentrations of Fe^2+^ were incubated with 100 µM of spermidine for 2 minutes. 100 µl of NBT (5 mg/ml) was added to the mixture and incubated at RT for another 5 minutes. The absorbance was recorded at 575 nm using UV-1800 Shimandzu UV-spectrophotometer.

### Quantitative real-time PCR (RT-qPCR)

Bacterial mRNAs were isolated by TRIzol^®^ reagent and the Qiagen bacterial RNA isolation Kit. DNase I treatment was done to remove residual DNA contaminant, and the integrity of the mRNA was checked on a 1% agarose gel. The RNA concentration was determined by a Nano-drop spectrophotometer (Thermo Scientific) and by a UV-1800 Shimandzu UV-spectrophotometer. 200ng of RNA samples, primer pairs (Table 2), and GoTaq^®^ 1-Step RT-qPCR System (Promega) were used for RT-qPCR. Reaction mixture without template were included as negative controls. At least three independent experiments were conducted for the determination of cycle threshold (C_T_) values. Fold expression change between spermidine-fed and unfed samples was calculated by the ΔΔC_T_ method. The values were normalized to the level of *betB* mRNA that was expressed constitutively as observed in the microarray.

### Other Biochemical assays

Different colorimetric assay kits (Sigma Aldrich) and ATP Bioluminescence assay Kit CLS II (Roche) were used to detect relative levels of NAD, NADH, GSH, and ATP. Aconitase assay was performed as per the protocol described (Gardner & Fridovich, 1992). Metal contents were determined by ICP-MS analyses at Punjab Biotechnology Incubator, Mohali, India. The metal concentration in the cell was determined as parts per billion (mg/kg) of *E. coli* cell pellets.

### Microarray experiments and interpretation

The saturated overnight culture of Δ*speG* strain was inoculated in the fresh LB medium and grown for 1.5 hours. After that 3.8 mM spermidine was added to one of the flasks, and the cultures were grown further for 2.5 hours. The cell pellets were harvested and washed with PBS, and dissolved in RLT buffer. The microarray was done from Genotypic Technology, Bangalore. The microarray had three probes for each gene on average.

### RNA Extraction and RNA Quality Control for Microarray

*E. coli* cell pellet was re-suspended in 300µl of 5mg/ml lysozyme and incubated at room temperature (RT) for 30 min. Isolation of RNA from *E. coli* was carried out using Qiagen RNeasy mini kit (Cat # 74106) as per manufacturer’s guidelines. A separate DNase treatment of the isolated total RNA was performed. The purity of the RNA was assessed using the Nanodrop Spectrophotometer (Thermo Scientific; ND-1000), and the integrity of the RNA was analyzed on the Bioanalyzer (Agilent 2100). We considered RNA to be of good quality based on the 260/280 values (Nanodrop), rRNA 28S/18S ratios, and RNA integrity number (RIN) (Bioanalyzer).

### Microarray Labeling

The sample labeling was performed using Quick-Amp Labeling Kit, One Color (Agilent Technologies, Part Number: 5190-0442). 500ng of each sample were denatured along with WT primer with a T7 polymerase promoter. The cDNA master mix was added to the denatured RNA sample and incubated at 40°C for 2 hours for double-stranded cDNA synthesis. Synthesized double-stranded cDNA was used as a template for cRNA generation. cRNA was generated by *in vitro* transcription, and the Cyanine-3-CTP (Cy3-CTP) dye incorporated during this step and incubated at 40°C for 2:30 hours. The Cy3-CTP labeled cRNA sample was purified using the Qiagen RNeasy column (Qiagen, Cat # 74106). The concentration of cRNA and dye incorporation was determined using Nanodrop-1000.

### Microarray hybridization and scanning

About 4 micrograms of labelled Cy-3-CTP cRNA was fragmented at 60°C for 30 minutes, and the reaction was stopped by adding 2X GE HI-RPM hybridization buffer (Agilent Technologies, In situ Hybridization kit, Part Number: 5190-0404). The hybridization was carried out in Agilent’s Surehyb Chambers at 65°C for 16 hours. The hybridized slides were washed using Gene Expression Wash Buffer 1 (Agilent Technologies, Part Number 5188-5325) and Gene Expression Wash Buffer 2 (Agilent Technologies, Part Number 5188-5326) and were scanned using Agilent Scanner (Agilent Technologies, Part Number G2600D). Data extraction from the images was done using Feature Extraction Software Version 11.5.1.1 of Agilent.

### Microarray data analysis

Microarray data analysis was undertaken by in-house coded R Script (https://cran.r-project.org/). Processing of raw data into expression profiles was achieved by utilizing the packages limma and affy. Probe intensities were converted into expression measures by standard procedures. Briefly, the design-sets depicting the “control/test” arrays were carefully generated by reading the raw data from MA image files. Background correction was done by the method “normexp”. This data was quantile normalized (between arrays depending on the design set), and within-array replicates were averaged. Processed data were categorized into major functional categories and tabulated. The detailed microarray array discussed in this manuscript have been deposited in GEO with accession number GSE154618.

## Acknowledgements

The authors are grateful to Dr. Debashish Adhikari, Division of Chemical Sciences, IISER Mohali, for their critical inputs on the plausible binding mechanism of spermidine and iron. The work has been funded by CSIR IMTECH, India to DD. VK was an ICMR fellow, RKM is a UGC fellow, DG is a CSIR fellow, AK is a DST-Inspire fellow, and AP is a DBT fellow.

## Author Contributions

VK, RKM, DG, AK and AP performed the experiments and discussed the data with DD. AA analyzed microarray gene expression raw data. GM analyzed data for ITC, bipyridyl and NBT assays to figure our coordination complex formation. AA and GM provided critical inputs to enhance the quality of work. DD conceptualized the work, analyzed the data and wrote the manuscript.

## Data Availability

All data are available in the main text, the supplementary files and source data.

---

**Figure 1- figure supplement 1.**
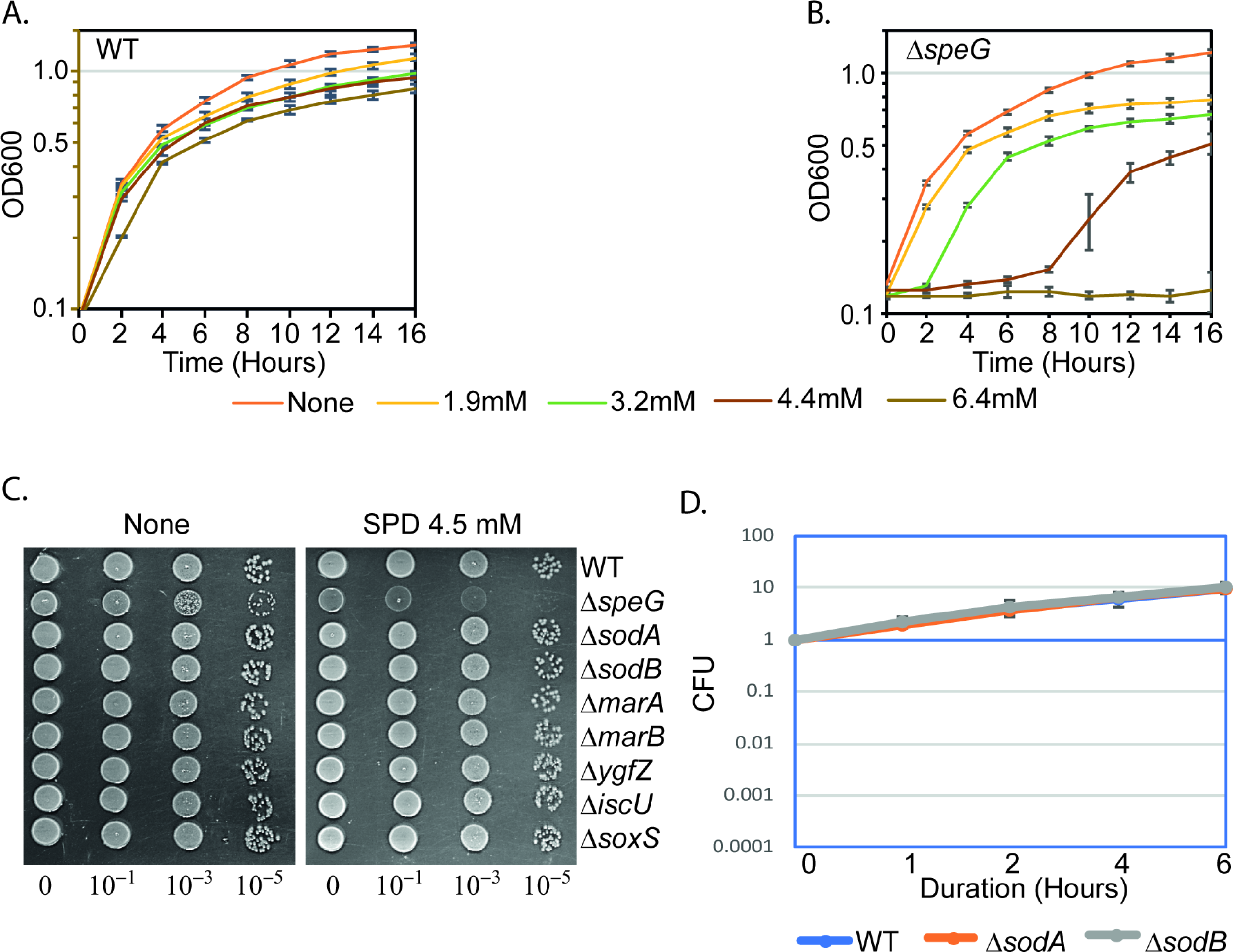
Spermidine-mediated O_2_^-^ production is toxic in the absence of SpeG function. **A** and **B** represents the growth curves of *E. coli* WT and Δ*speG* strains, respectively, in the presence of varying concentrations of spermidine. **C** and **D.** Spot and cell viability assays show that single Δ*sodA* and Δ*sodB* mutants are not affected by SPD. **Source data 1-** Figure 1A raw data **Source data 2-** Figure 1B raw data **Source data 3-** Figure 1C raw data **Source data 4-** Figure 1F raw data

**Figure 4 - figure supplement 1.**
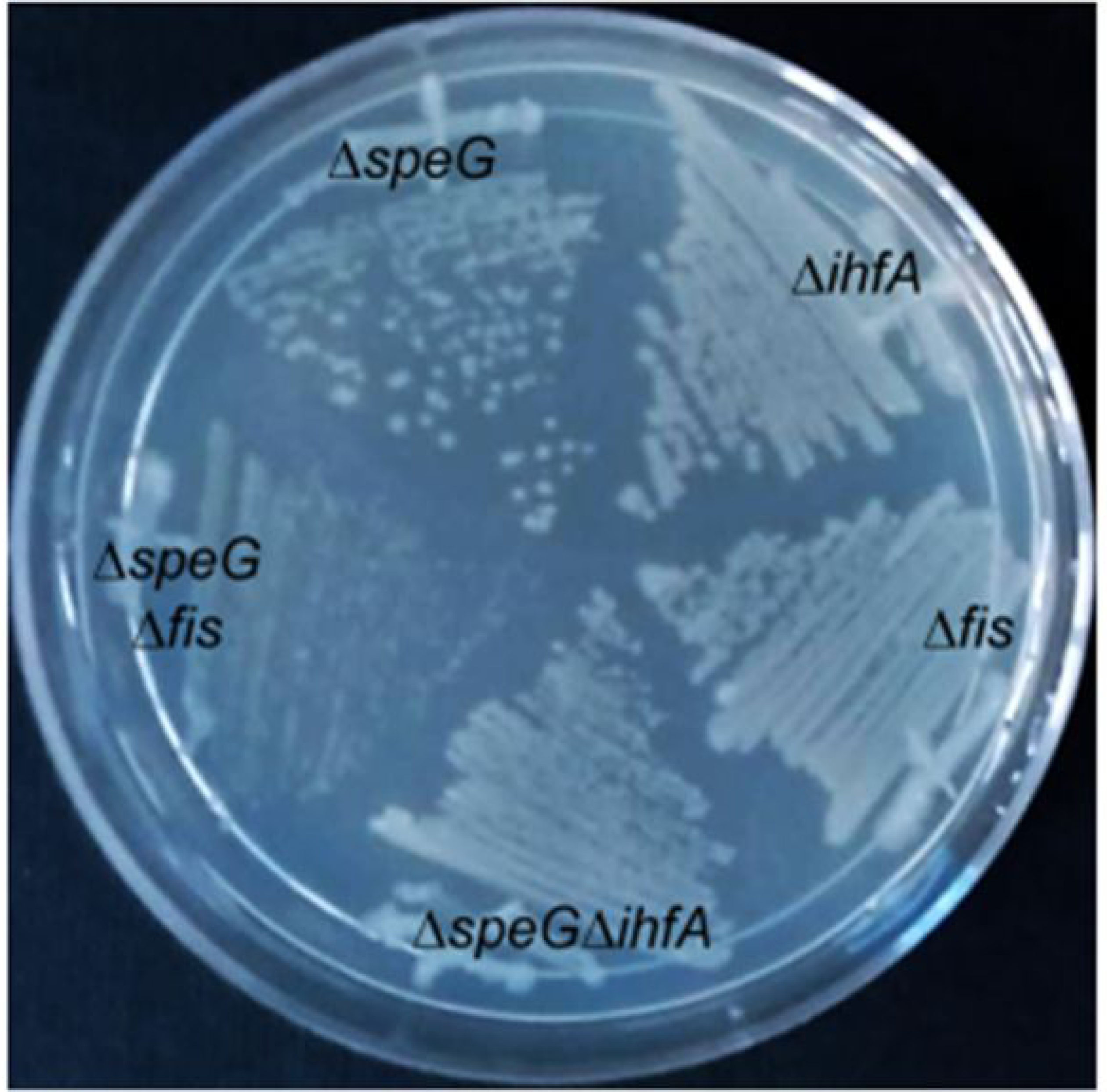
Indication of the importance of Fis regulator under spermidine stress. The colonies of different mutant *E. coli* strains were steaked on the LB agar surface to show that Δ*speG*Δ*fis* slow growing compared to Δ*speG* and Δ*speG*Δ*ihfA* strains.

**Figure 4 - figure supplement 2.**
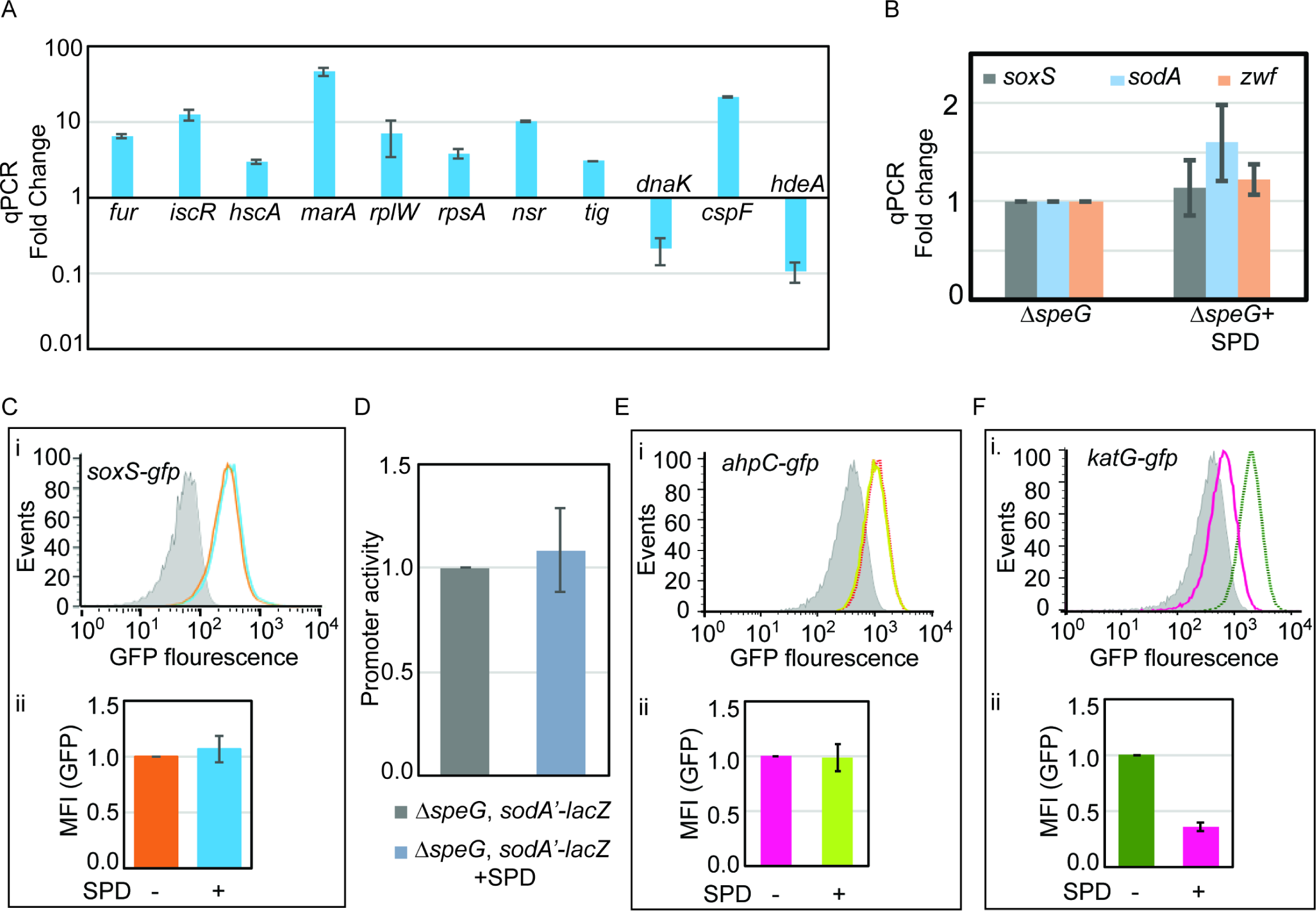
Validation of microarray data. A. Microarray data was validated by performing RT-qPCR against some of the genes that were up or down-regulated in the Microarray B. RT-qPCR data to show that the spermidine stress does not alter the expression of *soxS*, *sodA*, and *zwf* genes in the Δ*speG* strain. RT-qPCR data to show that the spermidine stress does not alter the expression of *soxS*, *sodA*, and *zwf* genes in the Δ*speG* strain. C. Flow cytometry experiments determined that spermidine stress does not upregulate P*soxS*- *gfpmut2* reporter in the Δ*speG* strain. The upper portion of the panel (i) represents a flow cytometry histogram. Grey area is background fluorescence of the Δ*speG* cells. The areas within saffron-line and blue-line represent GFP fluorescence in the absence and presence of spermidine, respectively. The bar diagram in the lower part of the panel (ii) exhibits relative MFI values calculated from three independent flow cytometry experiments. D. β-galactosidase reporter assay shows that the promoter activity of *sodA* gene remains unchanged under spermidine stress. E. Flow cytometry experiments were performed to show that spermidine stress does not upregulate P*ahpC-gfp* reporter in the Δ*speG* strain. The subpanel (i) represents a flow cytometry histogram of background fluorescence of the Δ*speG* cells (grey area). The pink-lined and green-lined areas of histogram represent fluorescence in the absence and presence of spermidine, respectively. The bar diagram in the subpanel (ii) exhibits relative MFI values calculated from three independent flow cytometry experiments. F. Flow cytometry experiments show that spermidine stress downregulates P*katG-gfp* reporter expression in the Δ*speG* strain. The subpanel (i) represents histogram of background fluorescence of the Δ*speG* cells (grey area). The green-lined and pink-lined areas of histogram represent GFP fluorescence in the absence and presence of spermidine, respectively. The bar diagram in the lower part of the panel (ii) exhibits relative MFI values in the presence or absence of spermidine calculated from three independent flow cytometry experiments. *** denotes P value <0.001; unpaired T-test. **Source data 12-** Figure 4B-ii raw data **Source data 13-** Figure 4C, 4D and 4E full images for western blots **Source data 14-** Figure 4C, 4D and 4E fold change values of the western blots **Source data 15-** Figure 4F raw data **Source data 16-** Figure 4G raw data

